# Divergence in the *Saccharomyces* species’ heat shock response is indicative of their thermal tolerance

**DOI:** 10.1101/2023.07.04.547718

**Authors:** Justin C. Fay, Javier Alonso-del-Real, James H. Miller, Amparo Querol

## Abstract

The *Saccharomyces* species have diverged in their thermal growth profile. Both *S. cerevisiae* and *S. paradoxus* grow at temperatures well above the maximum growth temperature of *S. kudriavzevii* and *S. uvarum*, but grow more poorly at lower temperatures. In response to thermal shifts, organisms activate a stress response that includes heat shock proteins involved in protein homeostasis and acquisition of thermal tolerance. To determine whether *Saccharomyces* species have diverged in their response to temperature we measured changes in gene expression in response to a 12°C increase or decrease in temperature for four *Saccharomyces* species and their six pairwise hybrids. To ensure coverage of subtelomeric gene families we sequenced, assembled and annotated a complete *S. uvarum* genome. All the strains exhibited a stronger response to heat than cold treatment. In response to heat, the cryophilic species showed a stronger response than the thermophilic species. The hybrids showed a mixture of parental stress responses depending on the time point. After the initial response, hybrids with a thermophilic parent were more similar to *S. cerevisiae* and *S. paradoxus*, and the *S. cerevisiae* x *S. paradoxus* hybrid showed the weakest heat shock response. Within the hybrids a small subset of temperature responsive genes showed species specific responses but most were also hybrid specific. Our results show that divergence in the heat shock response is indicative of a strain’s thermal tolerance, suggesting that cellular factors that signal heat stress or resolve heat induced changes are relevant to thermal divergence in the *Saccharomyces* species.

## Introduction

Temperature is a universal life parameter and organisms vary widely in their temperature tolerance. Temperature also varies over time and most organisms exhibit widespread changes in gene expression in response to temperature shifts (Lindquist 1986; Hofmann et al. 2000). In yeast, most temperature induced changes in expression are related to a general environmental stress response, characterized by largely transient changes in genes related to protein synthesis, cellular growth and metabolism (Gasch et al. 2000). In response to heat, cells also activate heat shock proteins, many of which are molecular chaperones that help disaggregate and refold denatured proteins (Richter et al. 2010). Whereas the heat shock response is present in most organisms, the threshold temperature for induction varies across organisms and is correlated with their optimal growth temperature (Lindquist 1986; Feder and Hofmann 1999). These and other observations implicate the heat shock response as a trait important to the ecology and evolution of most organisms (Tomanek 2010).

The heat shock response is an important mechanism of thermal tolerance (Richter et al. 2010). However, the fitness effects of the heat shock response are complex. In yeast, only a small fraction of proteins that are induced by heat are required to survival thermal stress (Gibney et al. 2013). The heat shock response also generates acquired thermotolerance whereby a mild heat stress increases survival of a later more severe heat stress. Acquired stress tolerance can also be activated by and protective against a variety of stresses besides heat (Berry et al. 2011). Both a cell’s stress response as well as its resistance to stress can depend on its growth rate, cell cycle phase and metabolic state (Lu et al. 2009; Slavov et al. 2012).

The role of the heat shock response in thermal adaptation within and between species is not as well characterized. In some cases heat shock proteins have been linked to thermal adaptation. After experimental evolution at high temperature, yeast strains were found to have constitutively activated heat shock proteins, even at low temperatures (Satomura et al. 2013; Caspeta et al. 2016). Expression of heat shock proteins has also been linked to thermal adaptation in a variety of other organisms but many of these observations are only correlative (Feder and Hofmann 1999).

The *Saccharomyces* yeast species exhibit substantial differences in their thermal growth profiles and provide an advantageous system to examine thermal adaptation (Gonçalves et al. 2011; Salvadó et al. 2011; Baker et al. 2019). Both *S. cerevisiae* and *S. paradoxus* can grow at 37°C, and at higher temperatures *S. cerevisiae* grows better than *S. paradoxus*. In comparison, *S. uvarum*, *S. eubayanus* and *S. kudriavzevii* cannot grow at 37°C but they grow faster than *S. cerevisiae* at low temperatures. These thermal growth differences impact competitive growth differences, and are particularly relevant to wine making (Williams et al. 2015; Alonso-del-Real et al. 2017). The two most thermally divergent species, *S. cerevisiae* and *S. uvarum*, also differ in their ability to survive heat shock treatment (Li 2018).

A number of studies have investigated the genetic basis of thermal divergence in the *Saccharomyces* species. Glycolytic enzymes have divergence in their activity and thermal stability (Gonçalves et al. 2011). The mitochondrial genome contributes to growth differences at high and low temperatures (Baker et al. 2019; Li et al. 2019). A genome wide screen identified eight genes involved in cell division and growth that provide *S. cerevisiae* with higher thermal tolerance relative to *S. paradoxus* but compromise growth at low temperature (Weiss et al. 2018; AlZaben et al. 2021). In *S. cerevisiae* and *S. uvarum* hybrids there are abundant *cis*-acting differences in gene expression but relatively few of these differences depend on the temperature at which the hybrid was grown (Li and Fay 2017).

Thermal growth differences among the *Saccharomyces* species are typically captured by monitoring growth after cells are shifted to a new temperature. The goal of this study was to examine the initial heat and cold shock response prior to any growth differences in order to determine whether and how these species have diverged in their temperature response. Species’ hybrids show that thermal tolerance is dominant, raising the question of whether thermally protective alleles mediate their effects prior to or subsequent to temperature changes and the heat shock response. Our results identify a small number of genes consistent with protective effects but show that most of the response to thermal treatment is indicative of a strain’s inherent thermal tolerance.

## Results

To examine divergence in heat and cold shock responses, gene expression was measured at four time-points (0, 15, 30 and 60 minutes) after a shift from 25°C to a low (12°C) or high (37°C) temperature for four *Saccharomyces* species and their six interspecific hybrids. Gene expression was measured using species-specific read counts from RNA sequencing and alignment to complete genomes of each species. Because only a draft genome with annotations was available for *S. uvarum*, we generated a complete chromosome-level assembly along with gene annotations (see Methods) and identified 5,121 genes with one-to-one orthologs in each of the four species.

### Thermal response in parental species

Across the four species more genes showed altered expression in response to heat (4,639) than to cold (1,874)(FDR < 0.01, Table 1), consistent with prior work in *S. cerevisiae* (Sahara et al. 2002; Schade et al. 2004). The average thermal response was similar in the two thermophilic species, *S. cerevisiae* and *S. paradoxus*, whereas it was initially smaller or slower in *S. kudriavzevii* in response to both temperature shifts, and stronger in *S. uvarum* in response to heat (Figure 1).

**Figure 1.**
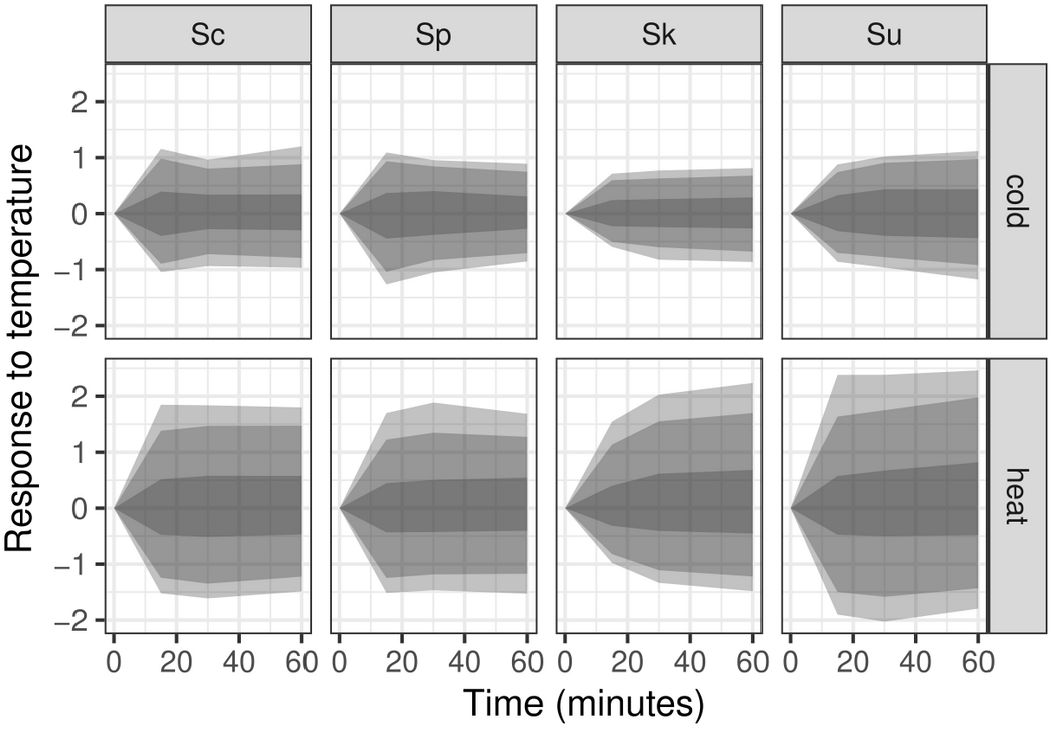
Species’ response to heat and cold treatment. Genes that significantly changed over time in response to heat or cold treatment were normalized to the zero time-point and shaded areas show the 50%, 90% and 95% quantiles (from darkest to lightest) of log2 changes in gene expression. Species are labeled with the first letter of their genus and species name: Sc, Sp, Sk and Su.

**Table 1.** Numbers of significant genes.

| Strain | Condition | Time | Allele | Time*Allele |
| --- | --- | --- | --- | --- |
| ScxSp | heat shock | 1911 | 745 | 12 |
| ScxSk | heat shock | 2303 | 1951 | 186 |
| ScxSu | heat shock | 2680 | 1724 | 100 |
| SpxSk | heat shock | 3630 | 2723 | 512 |
| SpxSu | heat shock | 3528 | 2871 | 417 |
| SkxSu | heat shock | 3183 | 2364 | 310 |
| Parents | heat shock | 4639 | 4827 | 3125 |
| ScxSp | cold shock | 417 | 988 | 0 |
| ScxSk | cold shock | 104 | 996 | 0 |
| ScxSu | cold shock | 757 | 1715 | 6 |
| SpxSk | cold shock | 1290 | 2467 | 13 |
| SpxSu | cold shock | 1105 | 2485 | 9 |
| SkxSu | cold shock | 1482 | 2669 | 38 |
| Parents | cold shock | 1874 | 4570 | 445 |

To examine species differences in their heat and cold shock response we used 3,125 heat responsive and 445 cold responsive genes with a significant interaction between the species’ alleles and time (FDR < 0.01, Table 1). Using expression values normalized to the initial time point, we identified genes with large expression differences between the two thermophilic species and the two cryophilic species. In response to heat, there were 197 genes whose average expression was 2-fold higher in the cryophilic compared to the thermophilic species. These genes were significantly enriched for involvement in *de novo* post-translational protein folding (n = 8, p = 7.0e-5, Table S1), and showed increased expression in response to heat and a larger increase in the cryophilic species (Figure 2A). There were 50 genes whose average expression was 2-fold lower in the cryophilic compared to thermophilic species. These genes were significantly enriched for involvement in cytoplasmic translation (n = 14, p = 3.5e-10, Table S1), and showed lower expression in response to heat in the cryophilic species (Figure 2B). In response to cold shock, there were only 11 genes with a 2-fold difference between the thermophilic and cryophilic species. Together, these results show that the cryophilic species exhibit a stronger heat shock response than the thermophilic species for a subset of genes.

**Figure 2.**
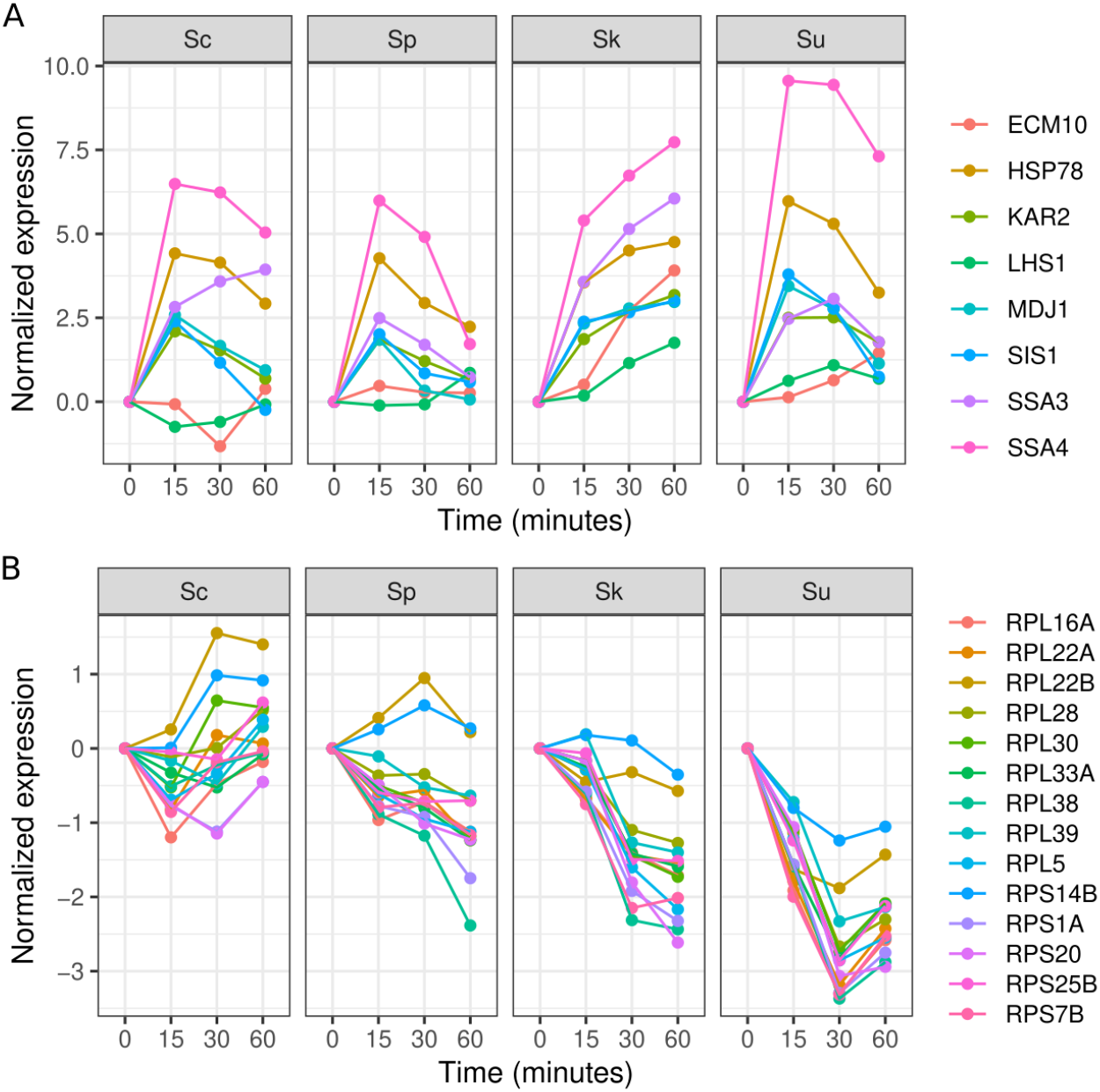
Genes that exhibit a stronger heat shock response in *S. kudriavzevii* and *S. uvarum* compared to *S. cerevisiae* and *S. paradoxus*. A. Eight genes involved in *de novo* post-translational protein folding with a 2-fold higher average response in the cryophilic species. B. Eleven genes involved in cytoplasmic translation with a 2-fold lower average response in the cryophilic species.

### Thermal response in species’ hybrids

Gene expression differences between species can be caused any number of cellular factors involved in the detection and response to thermal treatment. All the hybrids except *S. kudriavzevii* x *S. uvarum* grow at 37°C (Li et al. 2019). We thus examined whether hybrids with either an *S. cerevisiae* or *S. paradox* parent adopted a heat response similar to their parents.

The hybrids showed a large but variable number of genes that responded to temperature (FDR < 0.01, Table 1). Consistent with hybrid thermal growth profiles, hybrids with an *S. cerevisiae* parent had fewer genes that changed in response to heat or cold, and the *S. cerevisiae* x *S. paradoxus* hybrid had the fewest heat responsive genes of all the hybrids. To facilitate comparisons among hybrid gene expression, we used principle component analysis (PCA) of all genes regardless of which hybrids showed significant time dependent changes.

For the heat shock response, the first component explained 58% of the variance and the remaining components each explained less than 6% of the variance. The first component was associated most strongly with timepoint, followed by strain and allele (ANOVA R^2^ of 83%, 5.1% and 0.9%, respectively). Using the first component, we found the hybrids differed from one another and displayed a mixture of parental phenotypes that changed over time (Figure 3). At 15 minutes, *S. uvarum* showed the strongest response and *S. kudriavzevii* showed the weakest response among the parental species. At 15 minutes the hybrids were all more similar to *S. uvarum*, including the *S. cerevisiae* x *S. paradoxus* hybrid. At 30 minutes, the response of *S. kudriavzevii* and the *S. cerevisiae* x *S. kudriavzevii* hybrid was noticeably stronger than at 15 minutes. Except for the *S. cerevisiae* x *S. kudriavzevii* hybrid, all the hybrids with a thermophilic parent showed a reduced response at 30 compared to 15 minutes, and the *S. cerevisiae* and *S. paradoxus* hybrid response was weakest and similar to both of its parents. Finally, at 60 minutes most of the hybrids showed a reduced response, similar to that of *S. cerevisiae* and *S. paradoxus*. The two exceptions to this pattern were the *S. cerevisiae* x *S. kudriavzevii* and *S. kudriavzevii* x *S. uvarum* hybrids which retained their response level at or above *S. kudriavzevii* and *S. uvarum*.

**Figure 3.**
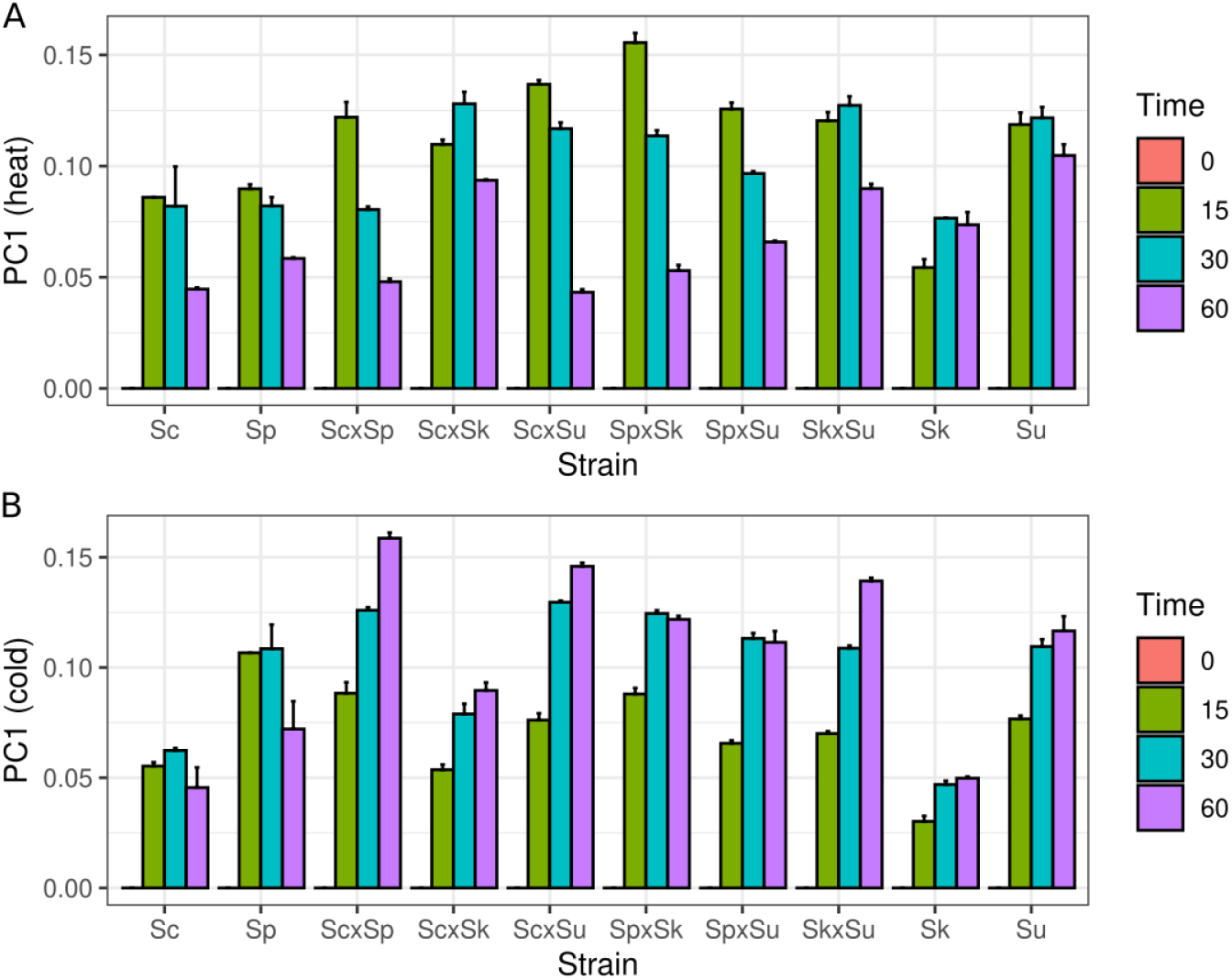
Hybrid and parental strains exhibit different heat and cold shock responses. Bars show the mean of the first principal component and whiskers show the standard error from replicates and species’ alleles for the heat shock (A) and cold shock (B) response. Because each gene’s expression was normalized to the zero time-point, the first component represents deviations from this time-point. Strains are represented by the first letter of the genus and species name: Sc, Sp, Sk and Su.

For the cold shock response, the first component accounted for 31% of the variance and was most strongly associated with time-point (ANOVA R^2^ = 77%). The overall pattern of the first component differed from the heat shock response (Figure 3). Like *S. kudriavzevii* and *S. uvarum*, all the hybrids showed an increasing response over time, with the 60 minute response being at or above the 30 minute response. Across all the time-points, the *S. cerevisiae* x *S. kudriavzevii* hybrid showed the weakest response, similar to its parental species. Considering the heat and cold responses together, the predominant pattern of gene expression as measured by PCA is that the hybrids differ from each other as well as from their parents and these differences change over time.

An alternative way of examining dominance in the hybrid is to focus on those genes that differed among the parental species. We thus considered genes that were 2-fold higher (197) or 2-fold lower (50) in the cryophilic species and examined their expression in the hybrids. Normalized to the initial time-point, the average heat response in the thermophilic species hybrid (ScxSp) and cryophilic species hybrid (SkxSu) mimicked differences found in their parents (Figure 4A). Genes that were 2-fold or higher in the cryophilic species (Sk and Su) were also higher in the cryophilic (SkxSu) compared to the thermophilic hybrid (ScxSp), particularly those induced by heat. Similarly, genes that were 2-fold lower in the cryophilic parents were also lower in the cryophilic hybrid. In comparison, these same genes did not mimic thermo/cryophilic differences in comparisons of hybrids containing one thermophilic and one cryophilic parent (ScxSk, SpxSk, ScxSu, SpxSu, Figure 4B and 4C). In comparison to the thermophilic (ScxSp) or cyrophilic (SkxSu) hybrid, these hybrids displayed intermediate phenotypes (Figure S1). These results show that similar to all genes, genes with a stronger heat shock response in the cryophilic species show the strongest response in the SkxSu hybrid and the weakest response in the ScxSp hybrid.

**Figure 4.**
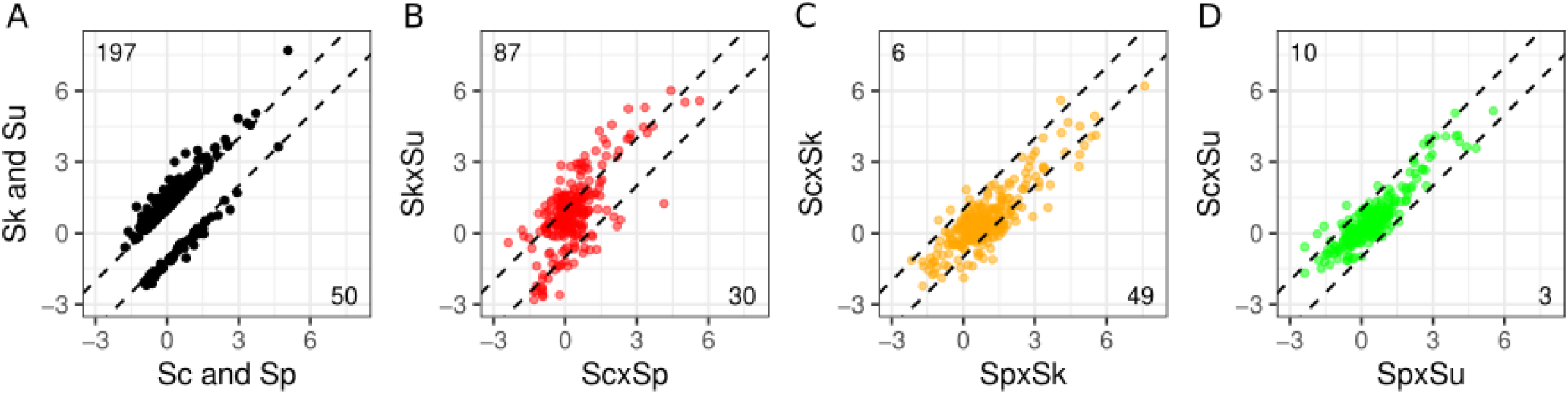
Hybrids recapitulate parental strain differences. Hybrid expression is shown in comparison to (A) genes that differed two fold between thermophilic (Sc and Sp, x-axis) and cryophilic (Sk and Su, y-axis) species (black points and dashed lines). For the same genes, the thermophilic hybrid (ScxSp) and cryophilic hybrid (SkxSu) recapitulate many of the parental differences (B: red), whereas the two *S. kudriavzevii* hybrids (C: orange) and the two *S. uvarum* hybrids (D: green) did not. Expression is the average response to heat across time-points and species’ alleles.

### Allele differences in species’ hybrids

Whereas each hybrid showed a different temperature response, within each hybrid both species’ alleles experience the same cellular environment and enable *cis*-acting regulatory divergence to be quantified. We examined allele-specific expression differences within each hybrid separately. Compared to their parents, the hybrids showed fewer allele differences that changed over time. There was an average of 256 heat responsive and 11 cold responsive genes that exhibited significant interactions between the species’ alleles and time (FDR < 0.01, Table 1). Consistent with the number of genes with time-dependent changes in expression, the three hybrids with an *S. cerevisiae* parent had fewer genes with significant allele-time interactions in response to both the heat and cold treatment (Table 1). Although the hybrid of the two thermophilic species, *S. cerevisiae* and *S. paradoxus*, showed the fewest allele-time interactions in response to heat, these two species are also the most closely related.

We next examined whether there was any consistent relationship between genes with allele differences and how they responded to heat (Figure S2). Only two of the hybrids (ScxSp and SkxSu) showed an association between allele differences and heat response based on the number of genes with significant allele and time effects compared to all other genes (Chi-squared test, Bonferroni corrected P < 0.05, numbers shown in Figure S2 quadrants). Four hybrids (all but ScxSp and ScxSk) also showed an association based on the number of genes with significant allele-time interactions. However, the major trend for this later association was that genes with allele-time interactions were more likely to increase in response to heat, and not that alleles of thermophilic species were expressed higher than those from cryophilic species for either heat induced or heat repressed genes.

We also assessed whether there were any genes with a significant allele-time interaction that showed a consistent difference between the alleles of the thermophilic and cryophilic species. Most of the genes were significant in only one of the hybrids (775/1095). Of the remaining, 208 were significant in at least two of the four hybrids between thermophilic and cryophilic species, and 5 genes were significant in all four of the hybrids (Figure 5: *TDH1*, *UTR4*, *MCR1*, *PER33* and *DAP1*). All of these five genes showed a consistent expression differences between the thermophilic and cryophilic species’ alleles.

**Figure 5.**
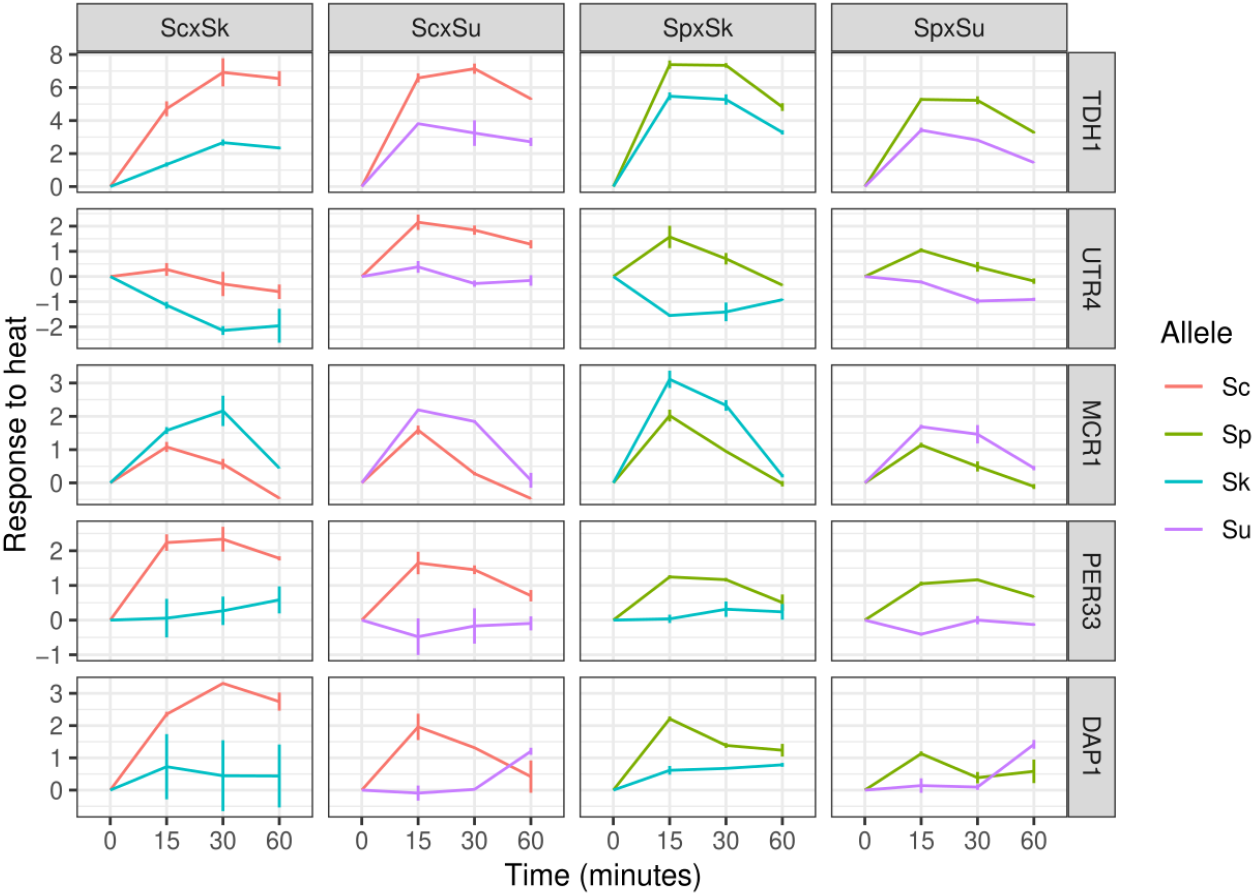
Five genes exhibit consistent allele differences between thermophilic and cryophilic species. Lines show the mean and whiskers show the point range.

### Thermal response of gene families

In addition to the 5,121 one-to-one orthologs, we also examined 321 gene families where the number of family members varied across species or where gene conversion between family members obfuscated one-to-one ortholog assignment. The gene families include many duplicated ribosomal genes, but also heat shock proteins (*HSC82*/*HSP82*, *SSB1*/*SSB2*, *HSP31-34*) and cell wall mannoproteins (*DAN*/*TIR*/*PAU*). To accommodate the variable number of family members we used the summed expression of all family members within a species. For these gene families, the fraction that showed time dependent and allele-time interactions was similar to that of one-to-one orthologs (Table S2). Only eight of these genes showed significant allele-time interactions between two of the four thermophilic-cryophilic hybrids. One of these families encodes 2-deoxyglucose-6-phosphate phosphatases (Randez-Gil et al. 1995), and has two copies in *S. cerevisiae* and *S. paradoxus* (DOG1/DOG2), but only one copy in *S. kudriavzevii* and *S. uvarum*. Whereas the *S. kudriavzevii* and *S. uvarum* alleles were strongly induced upon heat shock, the *S. cerevisiae* and *S. paradoxus* alleles were not (Figure S3).

## Discussion

The *Saccharomyces* species differ in their thermal growth profiles. These growth differences are rendered subsequent to any heat or cold shock response that occurs upon shift to a high or low temperature. Our study of gene expression in response to temperature shows that the *Saccharomyces* species as well as their hybrids differ substantially in their response to temperature, particularly heat. We find that major components of expression divergence are predicted by a strain’s thermal tolerance whereby sensitive strains exhibit the strongest heat shock response. Within the hybrids, we find abundant allele differences but relatively few that are consistent between alleles of thermophilic and cryophilic species. Together, our results suggest that thermal tolerance is present prior to or shortly after temperature exposure and generates many of the gene expression differences observed subsequent to temperature treatment.

Multiple aspects of the heat shock response are association with growth at high temperature. At 37°C, *S. kudriavzevii*, *S. uvarum* and their hybrid do not grow, but there is also variation in growth among the other strains (Li et al. 2019). *S. cerevisiae* and the ScxSp hybrid produce the largest colonies. The two other *S. cerevisiae* hybrids produce slightly smaller colonies, with ScxSu being larger than ScxSk. *S. paradoxus* colonies and its hybrids (SpxSk and SpxSu) are of equivalent size and smaller than those of *S. cerevisiae* and its hybrids. These growth differences mirror multiple aspects of the heat shock response. First, fewer genes showed significant responses to heat and allele interactions in hybrids with an *S. cerevisiae* parent, and the fewest in the ScxSp hybrid (Table 1). Second, the PCA analysis showed that the strength of the heat shock response at the later time-points corresponded to the heat sensitivity of the hybrids, with the ScxSk hybrid being one notable exception. Likewise, genes that were more strongly induced (protein folding) or repressed (ribosomal) in the cryophilic species were also more responsive to heat in the SkxSu compare to the ScxSp hybrid (Figure 4), and showed an intermediate response in the other hybrids (Figure S1).

Our findings are similar to a prior study of *Saccharomyces* hybrids that found a correspondence between heat shock response and growth at high temperature (Kempf et al. 2017). Using Hsp104-foci as a marker for protein aggregation, they found that an ScxSp hybrid resolved Hsp104-foci more rapidly than other *S. cerevisiae* hybrids. Additionally, they found that an ScxSk hybrid showed Hsp104-foci at a lower temperature and these foci reverted the slowest among all the hybrids.

In our study, interpreting expression of the ScxSk hybrid is not straightforward. The ScxSk hybrid, like its parents, is a uracil auxotroph. Although we did not observe substantial growth defects in Chardonnay grape juice, prior to heat shock the ScxSk hybrid and its parents showed higher expression of genes involved uracil biosynthesis (*URA1*, *URA2*, *URA8*) and lower expression of genes involved in lysine biosynthesis (*LYS4*, *LYS9*, *LYS12*, *LYS20*, *LYS21*). Equivocally, the hybrid also showed expression differences in glutathione metabolism (Figure S4). Glutathione metabolism is of interest since glutathione is a potent antioxidant and temperature causes oxidative stress (Morano et al. 2012), and secretion of glutathione has been shown to elevate the maximum temperature at which populations of *S. cerevisiae* can grow in a density dependent manner (Laman Trip and Youk 2020). In our experiments, glutathione synthetase (*GSH1*) was expressed at lower levels in *S. kudriavzevii* and the ScxSk hybrid compared to all other strains. The major glutathione-degrading enzyme *ECM38* showed higher expression of the *S. kudriavzevii* allele in both *S. kudriavzevii* and all of its hybrids. For both of these genes, the *S. cerevisiae* allele was more similar to *S. kudriavzevii* than *S. paradoxus*, making the ScxSk hybrid different from the SpxSk hybrid. Additionally, *GLR1*, which converts glutathione to its reduced form, showed greatly reduced expression in response to heat, but only in *S. cerevisiae* and for the *S. cerevisiae* allele in the ScxSk hybrid. The phospholipid hydroperoxide glutathione peroxidase (*GPX2*) uses glutathione to neutralize reactive oxygen species and showed low expression of the *S. kudriavzevii* allele in all the hybrids. Although uncertain as to the cause, the ScxSk hybrid showed both altered metabolism prior to heat shock as well as a heat shock response that stands out from the other hybrids.

We identified five genes that showed consistent allele differences between the thermophilic and cryophilic species. While none of them are directly linked to thermal tolerance, two are important for Erg11 activity and ergosterol biosynthesis. *MCR1* encodes a mitochondrial NADH-cytochrome b5 reductase that is one of two electron donors to the cytochrome P450 Erg11, and potentially Erg5 and Erg1 (Lamb et al. 1999). *DAP1* is a heme-binding protein that positively regulates Erg11 and Erg3 activity (Craven et al. 2007). In *S. cerevisiae*, inhibition of the later steps in ergosterol biosynthesis (Erg3-5) improves thermal tolerance, potentially through altered sterol composition of membrane lipids (Caspeta et al. 2014; Liu et al. 2017; Yang et al. 2023). However, the expression patterns of *MCR1* and *DAP1* may be incidental as they show opposite (compensatory) patterns in the thermophilic species.

Of the remaining three genes, *TDH1* functions in glycolysis and gluconeogenesis, *UTR4* is involved in the methionine salvage pathway, and *PER33* is associated with the ER and nuclear pore complexes. One interesting facet of *PER33* is that across many experimental conditions (SGD SPELL v2.0.3)(Hibbs et al. 2007) *PER33* expression is most correlated with *HCH1* (heat shock protein regulator), *CPR6* (chaperone activity with Hsp82), and *CUR1* (protein quality control); and its human homologue *TMEM33* is involved in the unfolded protein response (Sakabe et al. 2015).

Compared to heat shock, there were fewer genes that changed in response to cold treatment. This is consistent with prior work showing a small early response (<2hrs) and a larger later response (>2 hrs) that is related to the environmental stress response (Schade et al. 2004). We also found increased expression of genes previously described to be induced by cold treatment (*NSR1*, *TIP1*, *BFR2*, *OLE1*, *TIR1/2/4*). Similar to prior studies of log phase expression in hybrids (Li and Fay 2017; Hovhannisyan et al. 2020; Timouma et al. 2021), we found an appreciable number of allele differences, but very few allele differences that changed in response to cold treatment.

Relevant to both our heat and cold shock results, prior work has shown that gene expression in interspecific hybrids depends on which mitochondria they inherited (Solieri et al. 2008; Hewitt et al. 2020). The hybrids used in this study had mitochondria from their more thermophilic parent. The growth differences among hybrids depend on their mitochondria DNA and growth differences are mostly absent in petite hybrids lacking mitochondrial DNA (Li et al. 2019). It thus seems likely that the gene expression differences we observed may be quite different with mitochondria from the less thermophilic parent.

Gene families present a challenge to examining expression divergence when family members are closely related. Their analysis also depends on accurate annotation of complete genome sequences. For gene family members that couldn’t be assigned into one-to-one orthologous groups, e.g. varied in copy number among species, we used the sum of their expression values within each species. One interesting example is *DOG1/2*, which is present in two copies in *S. cerevisiae* and *S. paradoxus* but only a single copy in *S. kudriavzevii* and *S. uvarum*. The single copy *DOG1/2* alleles show a much stronger heat response. Mutations in *DOG1/2* confer 2-deoxyglucose resistance as Dog1/2 metabolizes derivatives of the glucose analog 2-deoxyglucose (Sanz et al. 1994). Although there are no links between *DOG1/2* and thermotolerance, the top *DOG1* phenome hit is sensitivity to fluvastatin, a statin that inhibits HMG-CoA reductase and ergosterol biosynthesis (Turco et al. 2023).

Among the other gene families, three are heat shock protein families. Whereas *SSB1/2* and *HSC82/HSP82* are present as closely related duplicates in each of the four species (consistent with duplication prior to speciation accompanied by gene conversion with each species), the *HSP31-34* family varies in copy number. *HSP31* is present in all four species as one-to-one orthologs, increases in response to heat, and shows much higher expression in the two thermophilic species (Figure S3). *HSP32* is only present in *S. cerevisiae*, *HSP33* is only present in *S. cerevisiae* and *S. paradoxus*, and *HSP34* is not present in any of the genome annotations. Members of the *HSP31* gene family have methylglyoxalase activity and redox-dependent chaperone activity that acts early in protein misfolding (Miller-Fleming et al. 2014; Amm et al. 2015; Tsai et al. 2015; Bankapalli et al. 2020). Deletions of *HSP31* family members cause reduced thermotolerance, sensitivity to oxidative stress, altered mitochondrial homeostasis and an increase in reactive oxygen species and oxidized gluthathione (Miller-Fleming et al. 2014; Bankapalli et al. 2020). These studies make the *HSP31* gene family a good candidate for functional studies of thermal divergence between species.

In summary, we find that thermal tolerance is indicative of the thermal response of the *Saccharomyces* species and their hybrids. What is the signal that turns on the heat shock response and why is it stronger in the thermally sensitive species? The heat shock transcription factor, *HSF1*, along with two stress response transcription factors, *MSN2* and *MSN4*, are responsible for most of the heat shock response (Morano et al. 2012). Phosphorylation of Msn2/4 by growth-related signaling pathways (cAMP/PKA) blocks their import to the nucleus and inhibits their activation of the stress response. Hsf1 is also post-translationally modified upon heat shock, and under the chaperone titration model, Hsf1 is negatively regulated by Hsp70, which is recruited away upon proteotoxic stress leading to Hsf1 activation of the heat shock response (Masser et al. 2020). However, heat shock also causes intracellular acidification and the formation of stress granules. Protein constituents of stress granules condense by phase separation in a temperature and pH dependent manner that may constitute an adaptive response to limit translation and protein misfolding (Glauninger et al. 2022). One such protein, Ded1, assembles into condensates in a temperature- and species-dependent manner such that the *S. kudriavzevii* protein does so at lower temperatures than that of *S. cerevisiae* (Iserman et al. 2020). Ded1 and other proteins involved in the translation mediated heat shock response condense at higher temperatures than those that activate Hsf1 but at lower temperatures than those which cause growth arrest. Thus, further characterization of the *Saccharomyces* species response to temperature could yield insight into the regulation of the heat shock response as well as to how it evolves.

## Material and methods

### Strains

A total of 6 diploid hybrid strains were made by crossing haploid strains of four *Saccharomyces* species (Table S3). The parental strains are derivatives of the following strains: T73 (*S. cerevisiae*), originally isolated from a vineyard in the Alicante wine region of Spain (Querol et al. 1992) and used as a commercial wine strain (Lallemand Inc., Montreal, Canada); N17 (*S. paradoxus*) originally isolated from oak exudate in Tartastan (Glushakova et al. 2007) and a member of the European population (Liti et al. 2009); CR85 (*S. kudriavzevii*) isolated from an oak tree bark in Agudo, Cuida Real, Spain (Lopes et al. 2010); BMV58 (*S. uvarum*) a commercial wine strain (Lallemand Inc.), originally isolated from wine in Utiel-Requena region of Spain (Henriques et al. 2021). Haploid derivatives of different mating types were obtained by replacing the *HO* gene with a dominant drug resistant marker (Goldstein and McCusker 1999; Vorachek-Warren and McCusker 2004), sporulation, and selecting haploid progeny by mating-type PCR (Huxley et al. 1990). Diploid hybrids with known mitochondrial types were generated by crossing parental strains with one parent lacking mtDNA (Li et al. 2019).

### RNA sequencing

Gene expression was measured by RNA sequencing of hybrids and their parental strains subjected to either a heat or cold shock. Two replicates of the six hybrids and the four parental strains were placed into 5 mL YPD (1% yeast extract, 2% peptone, 2% dextrose) and grown overnight at 25°C. Out of each of these pre-cultures, two sets (*A* and *B*) of four 2 mL tubes containing Chardonnay grape juice (Vintner’s Reserve Chardonnay, 10L, Cultures for Health) at 25°C were inoculated at an OD_600_ of 0.3, and placed at 25°C and 100 rpm for 5 hours. For thermal shock, sample sets *A* were placed at 12°C and sets *B* at 37°C. Sample collection for gene expression profiling was done at four different times: 0, 15, 30 and 60 minutes after thermal treatment. The total number of samples was 160: 10 strains, 4 time-points, 2 temperatures, 2 replicates. To reduce the number of sequencing libraries, cells from different tubes were mixed in order to bring the number of samples from 160 down to 64 (Table S4), with the criterion that none of the mixed strains shared the same genomic background (e.g. *S. cerevisiae x S. paradoxus* and *S. paradoxus x S. uvarum*). RNA samples were enriched for mRNA with the NEBNext Poly(A) mRNA Magnetic Isolation Module (NEB: New England Biolabs, Ispwich, MA); and library preparation was carried out by RNA fragmentation, cDNA synthesis, adaptor ligation, size selection and PCR enrichment using the NEBNext Ultra Directional RNA Library Prep Kit (NEB). PCR enrichment step was performed with NEBNext Multiplex Oligos for Illumina (Dual Index Primers, NEB #E7600), using 64 different combinations of indexed primers. A pool of the 64 libraries was run twice on a NextSeq Sequencing System from Illumina (2×75 bp run), generating a total of 936 million paired reads.

### Gene expression

For read mapping we selected representative genome assemblies from NCBI based on completeness and similarity to the strains used in this study. For *S. cerevisiae* we selected DBVPG6765 (GCA_002057805.1)(Yue et al. 2017), a European wine strain with Chr15R elongated by a 65-kb horizontal gene transfer from *Torulaspora microellipsoides* (Marsit et al. 2015). For *S. paradoxus*, we used CBS432 (GCA_002079055.1)(Yue et al. 2017), a common reference for European strains *(Liti et al. 2009)*. For *S. kudriavzevii* we used CR85 (GCA_003327635.1), and for *S. uvarum* we generated a genome assembly of YJF1449, a derivative of CBS7001 (described below). All assemblies contained complete chromosome level contigs with no gaps except for CR85, which had a six gene gaps between *PMT6* and *ADE3*, and an internal gap in *ATG13*, and DBVPG6765 and CBS432, which had their ribosomal RNA genes removed.

Reads were mapped to a combined reference of the four species’ genomes using TopHat (v2.1.1)(Kim et al. 2013) with a minimum and maximum intron length of 40 and 1500, respectively. HTSeq-Count (v0.9.1)(Anders et al. 2015) was used to count unique reads mapping to gene features using minimum alignment quality of 10. Over all the samples, 78% of read pairs were included yielding a median of 2.1 million counts per species-allele/sample with a range of 82 thousand to 16 million after removing one sample with very few reads (S26, *S. paradoxus*, cold, 15 min, Table S4). The median number of species-specific counts per gene was 125 for the 5,121 one-to-one orthologs (see Orthology assignments below).

### Genome sequencing

Two *S. uvarum* strains were used for genome assemblies: YJF1449 and YJF4719, both derivatives of CBS 7001 (Table S3). High molecular weight DNA was extracted (Barbitoff et al. 2021) with some modifications. Strains were grown in 500 mL YPD overnight at 25C in a shaking incubator at 300 rpm. After pelleting (2500 g for 5 min) and washing twice with DNAse-free distilled water, pellets were resuspended in 10 mL sorbitol lysis buffer (1M sorbitol, 0.05M EDTA, pH 8.0). Cells were treated with 100 uL zymolase (25 mg/mL, 20T) and incubated for 1 hour at 37°C to make spheroplasts. Spheroplasts were pelleted and resuspended in 10 mL total lysis buffer (0.1 M NaCL, 0.01 M Tris-HCl, 0.025 M EDTA, 0.5% w/v sodium dodecyl sulfate),and treated with 50 uL RNAse A (10 mg/mL) for two hours at 37°C. Lysates were treated with 100 uL proteinase K (20 mg/mL) overnight at 50°C. The next day, 15 mL 25:24:1 phenol:chloroform:isoamyl alcohol pH 8.0 (PCIaA) was added to each lysate and incubated for 10 minutes on a rotator. Lysates were added to phase separation tubes (50 mL conical tubes with 2 mL Dow Corning high vacuum grease pre-spun to the bottom of the tubes) and centrifuged at 4500 g for 5 minutes. The upper phase was transferred to new conical tubes where a second round of PCIaA extraction was performed. DNA was then precipitated with 1/10th volume 2M NaCl followed by 2/3 volume of isopropanol and mixing by gentle inversions after each addition. After a 10 minute incubation at room temperature, a glass hook was used to capture and place the DNA into 1.5 mL 70% ethanol, followed by 100% ethanol, and then dried. Precipitated DNA was rehydrated in 1 mL TE for 3 days. Size selection was performed using the Short Read Eliminator Kit (Circulomics).

Long read DNA libraries were prepared using the Ligation Sequencing Kit (Oxford Nanopore SQK-LSK110, version GDE_9109_v110_revJ_10Nov2020) without the addition of Lambda phage control DNA and were run through a R9.4.1 flow cell on a MinION. Each strain was sequenced on its own flow cell over which multiple rounds of sequencing libraries were run for 18-24 hours each. Between sequencing runs, the flow cell was cleaned and refreshed using the Flow Cell Wash Kit (EXP-WSH004, version SFC_9120_V1_revB_08Dec2020). Flow cells were reused until less than 5% of the pores were deemed available after washing. Base calling was performed using Guppy v.5.0.15 (Oxford Nanopore, dna_r9.4.1_450bps_sup.cfg configuration).

A second DNA extraction was performed for short read libraries using a YeaStar Genomic DNA Kit (Zymo Research). Libraries were generated using the Nextera DNA Flex Library Prep kit (Illumina) with unique dual indexes (IDT for Illumina) at 1/3 reaction volumes. Paired-end (2×100) sequencing was performed on a HiSeq (Illumina) by Novogene.

### Genome assembly

Short and long reads were filtered prior to assembly. Long reads were filtered for quality and length using Filtlong v.0.2.1 (https://github.com/rrwick/Filtlong), selecting for the highest quality reads at least 5000 bases long and targeting a 50X coverage of the estimated genome size (12 Mb). Short reads were trimmed and quality filtered using Trimmomatic v.0.36 (Bolger et al. 2014) with the following parameters: ILLUMINACLIP:nextera.fa:2:30:10:6 LEADING:20 TRAILING:20 SLIDINGWINDOW:4:15 MINLEN:50.

The filtered long reads were assembled with Canu v.2.2 (Koren et al. 2017), and then polished with all long reads that passed initial basecalling with Medaka v.1.5.0 (Oxford Nanopore), and then with 5 rounds of short read sequences using Polypolish v.0.5.0 (Wick and Holt 2022). The resulting assemblies were oriented and named according to the chromosome numbers assigned in the *S. uvarum* genome assembly for strain CBS 7001 (Chen et al. 2022)(NCBI accession number JAIPTR010000000). Two modifications were made to the YJF1449 assembly. One of the YJF1449 contigs contained a concatenated repeat of the complete mitochonrial sequence. This contig was oriented and trimmed to a single iteration by aligning to an existing *S. uvarum* mitochondion sequence (NCBI accession number KX657742). On chromosome XII, the YJF1449 assembly differed from that of YJF4719 and JAIPTR010000000 at the repeated rRNA locus. This region was thus reassembled and incorporated into YJF1449 chromosome XII using Gap5 v.1.2.14-r (Bonfield and Whitwham 2010). Reassembly was performed by extracting all Q10 and higher long reads in the region and assembling with Canu. Unlike the original assembly, the reassembled region showed even read coverage, similar to that obtained by mapping to the YJF4719 and JAIPTR010000000 assemblies and indicating that reassembly was more congruent with the input read data. The number of reads, bases and assembly statistics were tabulated for both strains (Table S5).

### Genome annotation

We generated genome annotations for each species. Our rationale was that we could use *S. cerevisiae* and *S. paradoxus* as controls to assess the quality and completeness of our annotation methods since they were previously accurately annotated using the LRSDAY pipeline (Yue et al. 2017; Yue and Liti 2018).

Each genome was soft masked with RepeatMasker (Smit et al. 2013) using the yeast transposon set (v1.0) from Casey Bergman (https://github.com/bergmanlab/yeast-transposons) and the ribosomal RNA genes (RDN5, RDN18, RDN58, RDN25) from S288C (SGD, R64-3-1). We used Augustus (Stanke et al. 2008) and YGAP (Proux-Wéra et al. 2012) to annotate protein coding genes (see below), tRNAscan-SE (v2.0.9)(Chan et al. 2021) for tRNA genes and MFannot (Prince et al. 2022) for mitochondrial genome annotations, available at https://megasun.bch.umontreal.ca/apps/mfannot/. LRSDAY also used tRNAscan-SE and MFannot and we found identical tRNA annotations for *S. cerevisiae* and *S. paradoxus* and nearly identical mitochondrial annotations (Table S6).

Augustus (v3.4.0) and YGAP were used to predict protein coding genes in the nuclear genome. We were unable to install and generate LRSDAY predictions for *S. kudriavzevii* and *S. uvarum*. Augustus was run with the saccharomyces_cerevisiae_S288C model, softmasking, hint parameters (extrinsic.M.RM.E.W.P.cfg), and with flags to report coding sequences with stop codons in gff3 format. We used RNA-seq alignments from TopHat (above), and scripts from Augustus to filter for unique hits (filterBam-uniq) and generated intron hints (bam2hints with max gene length = 10kb). A preliminary assessment of Augustus predictions using intron hints did not show increased concordance with LRSDAY annotations of DBVPG6765 and so we only used RNA-seq hints for manual curation and not for Augustus gene predictions. To generate protein hints, we used Spaln2 (v2.4.1)(Iwata and Gotoh 2012) to map *S. cerevisiae* proteins (5,943 genes annoated in S288c version R64-3-1 after excluding dubious genes) to each genome (max gene size = 10kb). Hits were filtered with the align2hints script from Augustus (max intron size 1500) and yielded hints for 5,909 *S. cerevisiae*, 5,756 *S. paradoxus*, 5,555 *S. kudriavzevii* and 5,471 *S. uvarum* proteins. For each predicted gene, IDs were designated by Augustus and a descriptive name was obtained by the best hit of the coding (DNA) sequence to all S288C coding genes using blastn (-word_size 9, -evalue 1e-40, -strand plus), or by the best blastp hit if no cDNA hit was returned. YGAP annotations were obtained online (Post-WGD species, No frame-shift correction). GenBank output files were converted to gff using bioperl’s bp_genbank2gff3 script.

### Comparison of gene predictions and manual curation

Augustus, YGAP, and LRSDAY annotations for *S. cerevisiae* and *S. paradoxus* were compared. Most annotations (86-90%) were exact matches across all three sets, with the same start/stop and intron/coding-exon positions (Table S6). Most of the remaining annotations were exact matches in two out of the three sets and involved slight differences in start/stop and intron positions. To assess the YGAP and Augustus annotations, we used manual curation based on protein and RNA-seq hints (described above) and gene structure in S288c (SGD, R64-3-1) including gene and intron size. The majority of annotations with exact matches in two of the three sets were curated as correct and we thus found clear mis-annotations in each set including missing introns and merged or split genes. In general, Augustus did better at annotating genes with introns and YGAP did better at annotating small genes. We only identified 17 *S. cerevisiae* and 31 *S. paradoxus* annotations that were curated as correct in LRSDAY but incorrect in both YGAP and Augustus (Table S6). Based on these results we curated all YGAP and Augustus annotations that did not match each other in *S. kudriavzevii* and *S. uvarum*. For each species, we generated a final set of annotations (Table S6) by combining our curation of nuclear proteins with annotations from tRNAscan and MFannot and repeats from RepeatMasker. For consistency in the gene expression analysis, we used these annotations rather than those from LRSDAY for *S. cerevisiae* and *S. paradoxus*.

### Orthology assignments

Hierarchical orthologous groups (HOGs) were defined by OrthoFinder (v2.5.4)(Emms and Kelly 2019). Hierarchical orthogroups are defined at each node in the species tree and OrthoFinder found 5,442 HOGs at the ancestral (N0) node, of which 5,270 had at least one ortholog in each species and 5,121 had one to one orthologs in all four species (Table S7).

### Expression analysis

Species-specific read counts were normalized using DESeq2 (Love et al. 2014). For each treatment (heat shock and cold shock) count data were fit to a model with interacting time and species-allele terms: counts ∼ time*allele. Genes with significant changes over time, between alleles or the interaction between the two were found by dropping those respective terms and evaluation using the negative binomial likelihood ratio test (nbinomLRT) with a false discovery rate cutoff of 0.01 (Benjamini and Hochberg 1995). For 321 gene families that did not have one to one orthologs in all four species we used the sum of the read counts across family members within each species.

For parental species differences, genes showing a significant interaction between allele and time were analyzed. For these genes, the average difference between the thermophilic and cryophilic species species was obtained from the species and time-point average after each gene was normalized to the initial time-point. GO term analysis was conducted using SGD with a background set equal to all 5,121 genes in the dataset and Holm-Bonferroni correction for multiple tests. Principal component analysis was used to compare the hybrid and parental species response to heat and cold treatment. Each gene’s expression was normalized to the zero time-point and principal components were obtained from the centered but unscaled expression data obtained from DESeq2’s variance stabilizing transformation function.

## Data availability

RNA sequencing reads are available at NCBI (PRJNA909640). DNA sequencing reads and assembly are available at NCBI (PRJNA882904). Processed data and analysis scripts are available at OSF (https://osf.io/7kunf/). These data are the genome annotations, read counts across the experimental conditions, normalized expression and p-values, and plots of gene expression data for each gene.

## Acknowledgments

This work was supported by a National Institutes of Health grant (GM080669) to JCF and the Spanish government MCIN/AEI to the Center of Excellence Accreditation Severo Ochoa (CEX2021-001189-S/funded by MCIN/AEI710.13039/501100011033).

