## Supporting Figures for "Divergence in the *Saccharomyces* species’ heat shock response is indicative of their thermal tolerance"

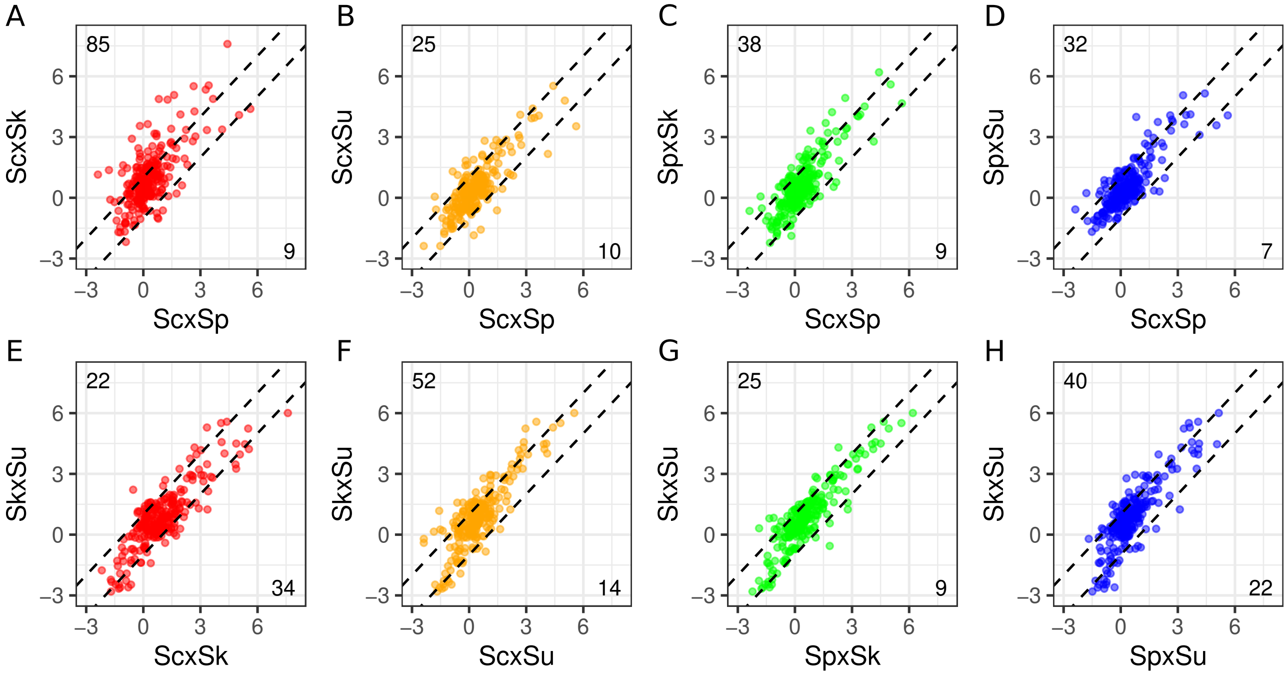


Figure S1. Hybrid expression differences in comparison to parental differences. Hybrid expression is shown in comparison to genes that differ two fold between thermophilic and cryophilic species (black dashed lines, see Figure 4). For the same genes, the ScxSp hybrid is shown in comparison to other hybrids (A: ScxSk, B: ScxSu, C: SpxSk, D: SpxSu) and the SkxSu hybrid is shown in comparison to other hybrids (E: ScxSk, F: ScxSu, G: SpxSk, H: SpxSu). Expression is the average response to heat across time-points and species' alleles.

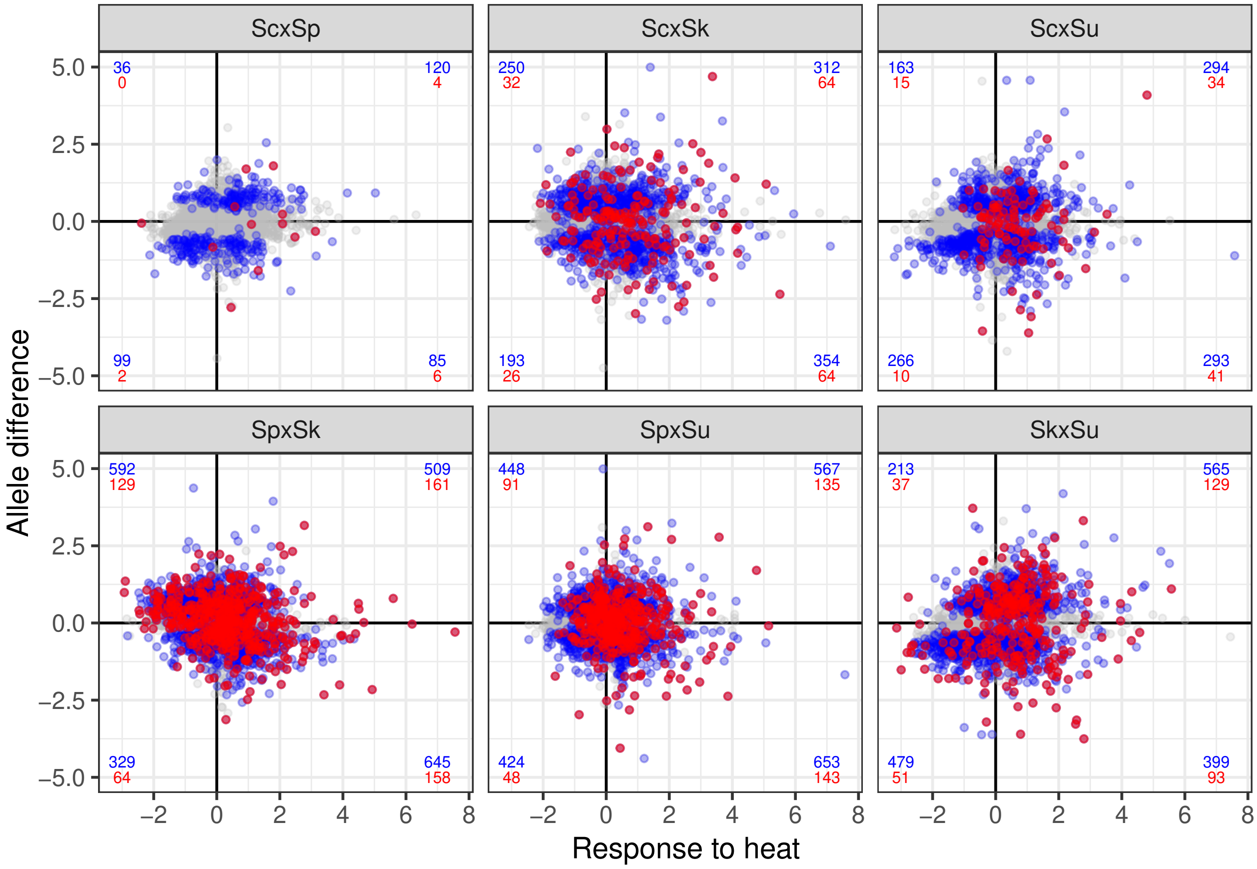


Figure S2. Relationship between allele differences and response to heat. The average allele difference across time-points (y, allele1/allele2 where alleles come from the hybrid name: allele1 x allele2) versus the average response to heat (x, normalized to the zero time point). Points show genes with significant allele and time effects (blue), genes with significant allele-time interactions (red), and all others (grey). Numbers in each quadrant indicate the number of genes with higher/lower expression for each species' allele and with an increase/decrease in response to heat, colored by significance category.

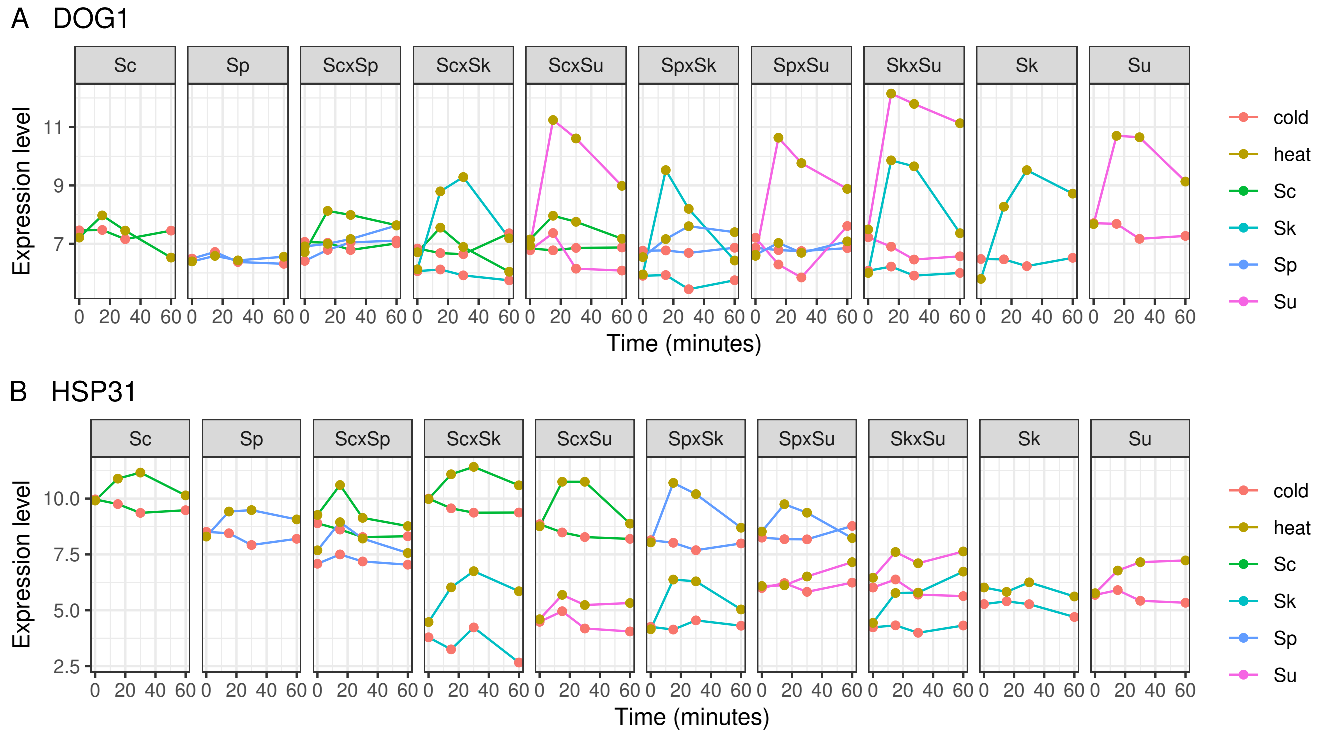


Figure S3. Expression of gene families in response to heat and cold treatment. For the DOG1 family (A), S. cerevisiae and S. paradoxus expression is the sum of DOG1 and DOG2, whereas S. kudriavzevii and S. uvarum expression is for a single copy. For HSP31 (B) only HSP31 expression is shown since one-to-one orthologs could be distinguished from other family members (HSP32-34).

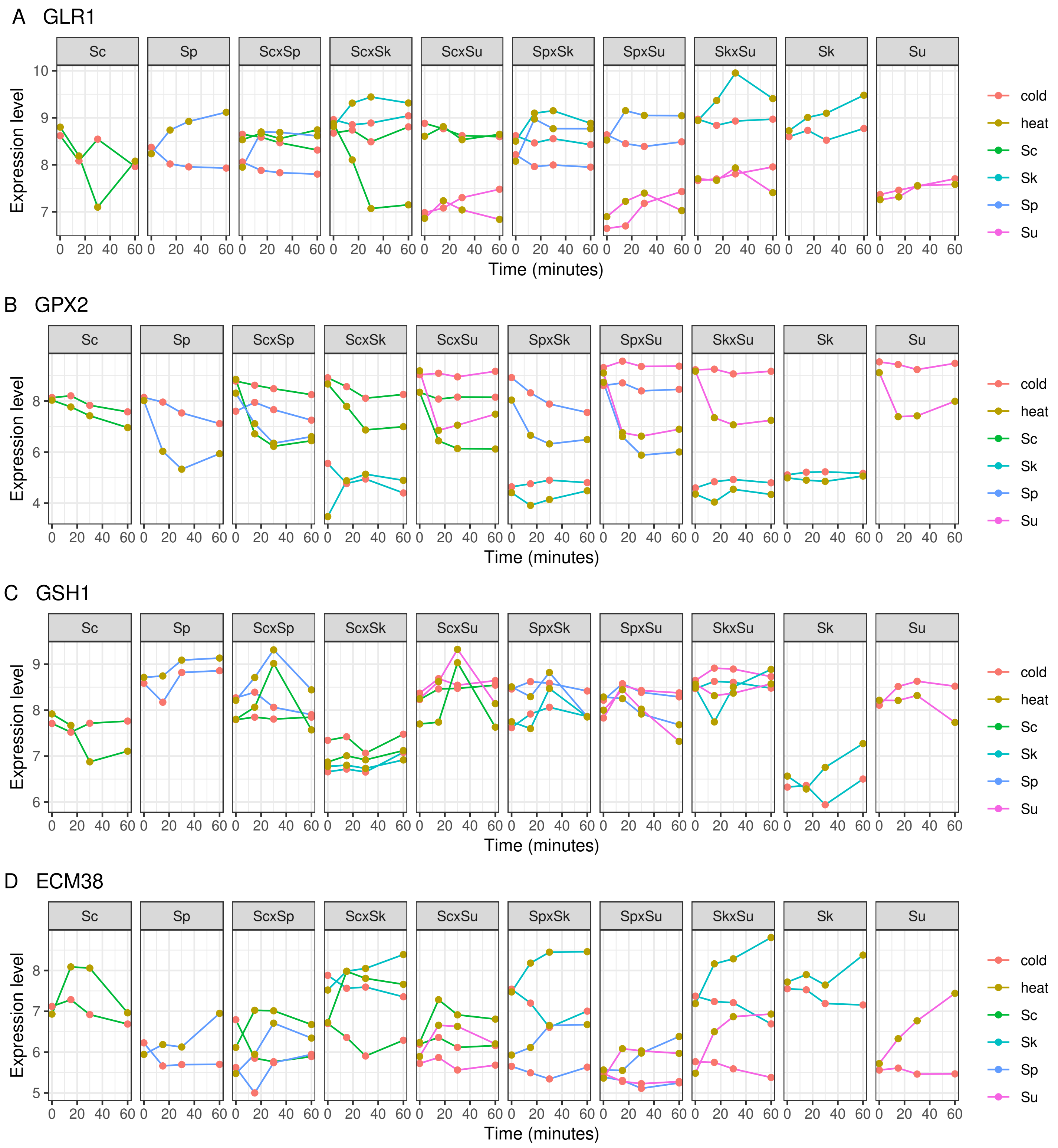


Figure S4. Expression of genes involved in glutathione metabolism: GLR1 (A), GPX2 (B), GSH1 (C) and ECM38 (D).
